# Effects of sexual dimorphism and estrous cycle on murine *Clostridioides difficile* infection

**DOI:** 10.1101/2023.07.05.547871

**Authors:** Jacqueline R. Phan, Erica Bacab, McKenzie Washington, Maria Niamba, Dung M. Do, Tiffany V. Mata, Amber Consul, Liahm Blank, Stephanie Yang, Amber J. Howerton, Efren Heredia, Robert Soriano, Chandler Hassan, Chad L. Cross, Ernesto Abel-Santos

## Abstract

*Clostridioides difficile* infection (CDI) is responsible for the majority of identifiable antibiotic-associated diarrhea. Women are more susceptible to CDI than men. In this study, we show that female mice developed more severe CDI than males. Furthermore, females in estrus developed only mild CDI 1-2 days later, while females in proestrus developed deadly disease. Mirroring the delayed effect of the estrous cycle, pre-infection prolactin levels formed a complex network with immunoglobulins and cytokines that affected early CDI severity one day after challenge. Similarly, pre-infection progesterone and luteinizing hormone formed a network that affected CDI two days after challenge. As expected, immune effectors early in the infection formed a hormone-independent network that concurrently correlated with CDI severity. Interestingly, early infection follicular stimulating hormone levels created a network that affected the CDI recovery phase. In summary, murine sexual hormones affect CDI progression by affecting the immune system both before and during disease progression.

## Background

*Clostridioides difficile* infection (CDI) is responsible for approximately 25% of all antibiotic-associated diarrhea [1, 2]. In the U.S., over 223,900 CDI cases occur annually, with a mortality rate of up to 6.5% with costs estimated to be $6.3 billion [3–5].

When adjusted for age, immune status, and antibiotic exposure, sex is also an important risk factor in human CDI. Contrary to other GI bacterial infections [6], women are more vulnerable than men to both primary CDI and CDI relapse [7–14]. Furthermore, CDI has been flagged as an emerging threat to pregnant and peripartum women [15–18]. Similarly, older women might be at higher risk of developing CDI due to permanent hormonal depletion after menopause.

An obvious factor underlying sexual dimorphism is the fluctuating hormone concentrations during the female ovulation cycle [19]. Sex hormones have strong effects on both human and murine metabolism [20] and can affect the immune and humoral responses [21–23], as well as susceptibility to bacterial infections [6]. Changes in sex hormone levels can also affect the gut microbiota [24–26], the main protective barrier against *C. difficile* colonization [27].

Similar to humans, female mice undergo hormonal changes during the estrous cycle [28]. Estradiol levels increase during the proestrus stage, followed by spikes in luteinizing hormone (LH) and follicular stimulating hormone (FSH) [29]. The estrus stage is dominated by rapid increase of prolactin (PRL), followed by a sharp decrease to trigger metestrus entry. The metestrus stage shows a sharp increase of progesterone, with a smaller simultaneous increase in estradiol. Finally, progesterone levels peak during the diestrus stage. Testosterone is produced uniformly throughout the estrous cycle, but its circulating concentration is 10-fold lower than in males [30]. Hence, each murine estrous cycle stage is characterized by a unique pattern of hormone levels.

We have shown that steroidal sex hormones can interact with *C. difficile* spores and modulate germination [31, 32]. Similarly, since spore germination is a necessary step for disease onset, we have found that sterane analogs can inhibit this process and prevent CDI onset in rodent models of disease [33–43]. During our search for CDI prophylactics, we serendipitously observed that male mice tended to develop less severe CDI than their female counterparts.

In this study, we used a murine model to assess for sexual dimorphism during CDI. We found that female mice developed more severe CDI signs than males when infected with three different *C. difficile* strains. Sexual dimorphism did not disappear under nutritional conditions known to trigger lethal murine CDI [44]. Within a normal infection, CDI sign severity in female mice correlated with the estrous stages 1-2 days prior to maximal CDI scores. As a result of this delayed effect, animals were mostly protected from CDI if they were in estrus 1-2 days before reaching maximal signs. In contrast, animals developed deadly disease signs 1-2 days after proestrus. Mirroring the delayed effect of the estrous cycle on CDI, we found that pre-infection levels of PRL, immunoglobulin IgG2b, cytokine IL-1β, cytokine G-CSF, and chemokine KC were the primary nodules of a complex network that correlated with CDI symptomatology on the day after infection. Similarly, we found that the pre-infection levels of progesterone, LH, immunoglobulin IgG1, chemokine eotaxin, and chemokine IP-10 were the primary network nodules that affected CDI severity but with a 2-day delay. As expected, the levels of immunoglobulins, cytokines, and chemokines during early infection formed a hormone-independent network that concurrently correlated with CDI severity. Interestingly, early infection levels of FSH, together with cytokine IL-1β, and chemokine KC were the main nodules of a network that affect CDI severity 2-days later, during the recovery phase of the infection. In summary, we show that murine female sexual hormones affect CDI progression, likely as a function of immune system modulation both before infection and during disease progression.

## Materials and Methods

### Materials

*C. difficile* strains R20291 and 9001966 were donated by Professor Nigel Minton at the University of Nottingham, United Kingdom. *C. difficile* strain 630 was obtained from the American Type Culture Collection (ATCC, Manassas, VA). The synthetic bile salt analogs, CAMSA and CAPA were prepared by Professor Steven M. Firestine at Wayne State University in Detroit, Michigan. Standard Laboratory Rodent Diet was obtained from LabDiet (St. Louis, MO, USA). Custom purified diet (Atkins-type diet) containing approximately 50% protein and 25% fast was purchased from TestDiet (TestDiet Catalog # 18181230286). All *C. difficile* strains were grown and maintained in a Coy Laboratories (Grass Lake, MI) vinyl anaerobic chamber (5% H2, 5% CO2, 95% N2).

### Animals

All procedures involving animals in this study were performed in accordance with the Guide for Care and Use of Laboratory Animals outlined by the National Institutes of Health. The protocol was reviewed and approved by the Institutional Animal Care and Use Committee (IACUC) at the University of Nevada, Las Vegas (Permit Number: R0914-297).

Weaned C57BL/6 mice were purchased from Charles River Laboratories (Wilmington, MA, USA). Mice were housed in groups of five per cage at the University of Nevada, Las Vegas animal care facility. Upon arrival at the facility, animals were allowed to acclimate for at least one week prior to the start of experimentation. All post-challenge manipulations were performed within a biosafety level 2 laminar flow hood.

### C. difficile spore harvest and purification

*C. difficile* cells from frozen stocks were streak plated in an anaerobic chamber onto BHI agar supplemented with 2% yeast extract, 0.1% L-cysteine-HCl, and 0.05% sodium taurocholate (BHIS) to yield colonies (15). After 48 hours, a single colony was inoculated into BHI broth supplemented with 0.5% yeast extract and incubated for 48 hours. The inoculated broth was then spread plated onto multiple BHI agar plates prepared as described above. Inoculated plates were incubated for 7 days at 37 °C.

Plates were then flooded with ice-cold deionized (DI) water. Cells and spores were harvested by scraping bacterial colonies from the plates. The cell and spore mixtures were pelleted via centrifugation at 8,000 ×g for 5 minutes, resuspended in DI water, and pelleted again. This washing step was repeated twice more. The cell/spore mixture was then centrifuged through a 20% to 50% HistoDenz gradient at 18,200 ×g for 30 minutes with no brake (22).

Under these conditions, spores pelleted at the bottom of the centrifuge tube, while cell debris remains above the 20% HistoDenz layer. Pelleted spores were transferred to a clean centrifuge tube and washed five times before storing in DI water at 4 °C. To determine spore purity, selected samples were stained using the Schaeffer-Fulton endospore staining method (23) or were visualized via phase-contrast microscopy. Spore preparations used were greater than 95% pure. Spores were washed three times with DI water, heat shocked at 68°C for 30 minutes, then washed three additional times before being resuspended in DI water prior to administration.

### Estrous cycle tracking in mice

The estrous cycle tracking method performed in this study was adapted from *McLean et al., 2012* [29]. Mice estrous stages were tracked for at least one week prior to any manipulation to ensure that animals were cycling normally. Animals were tracked daily for their estrous cycle stage via vaginal lavage at the same time each morning throughout the duration of the study. First, excess urine is dabbed off with low-lint Kimwipes. A micropipette was used to lavage 20 μL of PBS into the vaginal orifice of the animal. Water was swooshed 3-4 times around the vaginal orifice until the suspension appeared mildly cloudy. The 20 μL sample was then transferred onto a clean glass microscope slide and left to air dry for at least 1 hour. Animals were monitored to ensure there were no signs of vaginal infection.

Air-dried vaginal sample slides were dipped into a 2% crystal violet solution inside a Coplin jar for 1 minute. Slides were then moved to a Coplin jar containing fresh DI water to wash off excess stain. Slides were dabbed gently with clean wipes, allowed to air dry, and examined using a compound light microscope at 100x and 400x magnification (**Figure S1**) [45].

### Murine CDI model

The murine CDI model used in this study was adapted from Chen *et al.* [42]. Mice were given three consecutive days of antibiotic cocktail containing kanamycin (0.4 mg/ml), gentamycin (0.035 mg/ml), colistin (850 U/ml), metronidazole (0.215 mg/ml), and vancomycin (0.045 mg/ml) *ad libitum* (14). Mice were then given DI water for the remainder of the experiment. On the day prior to *C. difficile* challenge (day −1), mice were given an intraperitoneal (IP) injection of 10 mg/kg clindamycin. On the day of infection (day 0), experimental groups were challenged with 10^8^ *C. difficile* spores via oral gavage.

Mice were observed for signs of CDI twice daily and disease severity was scored according to a CDI sign rubric adapted from previously published work [36]. According to the rubric, pink anogenital area, mild wet tail, and weight loss of 8-15% were each given a score of 1. Red anogenital area, lethargy/distress, increased diarrhea/soiled bedding, hunched posture, and weight loss greater than 15% were each given a score of 2. Animals with a cumulative scored less than 3 were indistinguishable from noninfected controls and were considered non-diseased. Animals scoring 3–4 were considered to have mild CDI. Animals scoring 5–6 were considered to have moderate CDI. Animals scoring greater than 6 were considered to have severe CDI and were immediately culled.

### Tracking the estrous cycle in the murine CDI model

To better visualize the diverse range of murine CDI signs [46], the hypervirulent strain R20291 was employed for all estrous-related studies. Estrous stages for each mouse were tracked every morning at the same time via vaginal lavage and cytology. Mice that were not cycling or cycling irregularly were excluded from the study. In the first round of the study (cohort 1), animals were sacrificed at random on day 0 (prior to *C. difficile* challenge), day 1, day 2, and day 3 (post-*C. difficile* challenge) after daily estrous stage sampling. Animals that were severely ill were immediately culled regardless of day or estrous stage. Surviving animals on each day were analyzed for estrous stage and CDI signs.

After initial analysis of data from the first experiments showed that estrous stages have a delayed effect on CDI severity, the protocol was further modified in an attempt to specifically track certain estrous stages and collect non-terminal blood samples on the days prior to expected maximal CDI signs. In the second round (cohort 2), mice were pooled based on their predicted estrous stage on the day of spore challenge. This prediction was estimated from the estrous stage they were on at the start of antibiotic treatment five days prior to infection. For each estrous stage grouping, animals were randomly selected for blood sampling via facial vein bleed on day 0 (day of challenge) or day 1 (24 hours after challenge). Animals were monitored for CDI signs, as above and sacrificed either one day or two days after blood samples were collected.

### Blood sample processing for sex hormones, cytokines/chemokines, and immunoglobulins

Approximately 100 μl of blood was collected from each mouse via facial vein bleeds. The blood rested at room temperature for 30 minutes prior to spinning down to collect serum. Serum was frozen at −20°C and then stored at −80°C until processing. Each serum sample was divided into four aliquots.

The first aliquot was analyzed to determine the concentration of follicle-stimulating hormone (FSH), luteinizing hormone (LH), and prolactin (PRL) using the Milliplex Mouse Pituitary Magnetic Bean Panel from Millipore according to kit instructions on a Bio-Plex 200 system. The second aliquot was processed using Millipore’s Milliplex Mouse Cytokine/Chemokine Magnetic Bead Panel according to kit instructions to test for cytokines IL-13, IL-10, IL-2, IP-10, IL-5, RANTES, KC, IL-15, IL-1α, IL-1β, IL-17, IL-6, TNF-α, G-CSF, eotaxin, IFN-γ, and GM-CSF. The third aliquot was processed using Milliplex Mouse Immunoglobulin Isotyping Magnetic Bead Panel from Millipore according to kit instructions to test for IgM, IgG, IgA, IgE, and IgD.

The fourth aliquot was sent to The Metabolomics Innovation Centre (TMIC, Alberta, Canada) for analysis of progesterone concentrations by HPLC-MS. Briefly, 10 μl of plasma was extracted with 40 μl of acetonitrile containing 3.1 nM progesterone-d9 internal standard (Cayman chemical, Ann Arbor, MI). The extraction mixture was sonicated in a water bath for 10 minutes, centrifuged at 21000 x g for one minute, and 40 μl of the supernatant was transferred to the autosampler vials with glass inserts. LC-MRM-MS analysis was performed using a Nexera XR UPLC system (Shimadzu, Japan) coupled to a Sciex QTrap 6500+ mass spectrometer equipped with an IonDrive Turbo V electrospray ionization source. Chromatography was performed using a Zorbax Eclipse Plus C18 column (2.1 × 150 mm, 1.8 μm, Agilent, USA) at a flow rate of 0.2 mL/min of mobile phases 0.1% formic acid in water (A) and 0.1% formic acid in acetonitrile (B). The LC gradient was 50-100% B / 0.0-5.0 min, 100-50% B / 5.0-5.1 min, 50% B / 5.1-8.0 min. The sample injection volume was 10 μl, the column oven was off, and the autosampler temperature was 15°C. MRM-MS was performed using electrospray ionization in the positive mode. 315.2/97.1 and 324.3/100.1 Q1/Q3 transitions were monitored for progesterone and progesterone-d9, respectively (1). Dwell time was 50 ms, collision energy (CE) 32 V, declustering potential (DP) 50 V, entrance potential (EP) 10 V, collision cell exit potential (CXP) 12 V, source temperature 450 °C; ion source gas 1 40 psi; ion source gas 2 40 psi. The collision gas was set to high, and the curtain gas was set to 40 psi. Data acquisition and analysis was performed using Analyst 1.6.3 software (Sciex, Framingham, MA). Progesterone concentrations were determined from chromatographic peak area ratios of the analyte and the internal standard.

### Statistical analyses

For the murine CDI model, severity of signs was visualized via box-and-whisker plots with a minimum of three independent values (n ≥ 3). One-way ANOVA or one-tailed *t*-test were performed, as appropriate, using R (v. 4.4.3, R Core Team, 2025) to assess differences between groups at every time point, with significance defined as p < 0.05. Significant main-effect ANOVAs were analyzed using Scheffe’s correction for *post hoc* comparisons between all groups. Time-to-event data were analyzed using Kaplan-Meier survival analysis models, with log-rank tests for curve comparisons. For assessment of associations among variables, we utilized the robust Kendall’s Tau correlation coefficient for bivariate comparisons among hormone levels, cytokine levels, immunoglobulin levels, and CDI severity scores, with significance defined as p < 0.10. Correlations were also calculated for combined samples and individually by bleed day (day 0 or day 1).

## Results

### Female mice exhibit more severe CDI than male mice

We infected male and female C57BL/6 mice with three different *C. difficile* strains [33]. One group of male and female mice was infected with spores from *C. difficile* strain 9001966 (ribotype 106) (**Fig. 1A**). For both sexes, mice started to develop signs of CDI on day 2 post-challenge (day 2). Male mice showed varied CDI sign severities during days 2-4, but all animals recovered by day 5. Female mice, on the other hand, showed more homogenous mild to moderate CDI symptomatology in days 2-3 and all animals recovered by day 4. Even though the CDI timeline was shorter in female mice, these animals developed statistically more severe signs than their male counterpart at the peak of infection (**Fig. 1A**).

**Figure 1.**
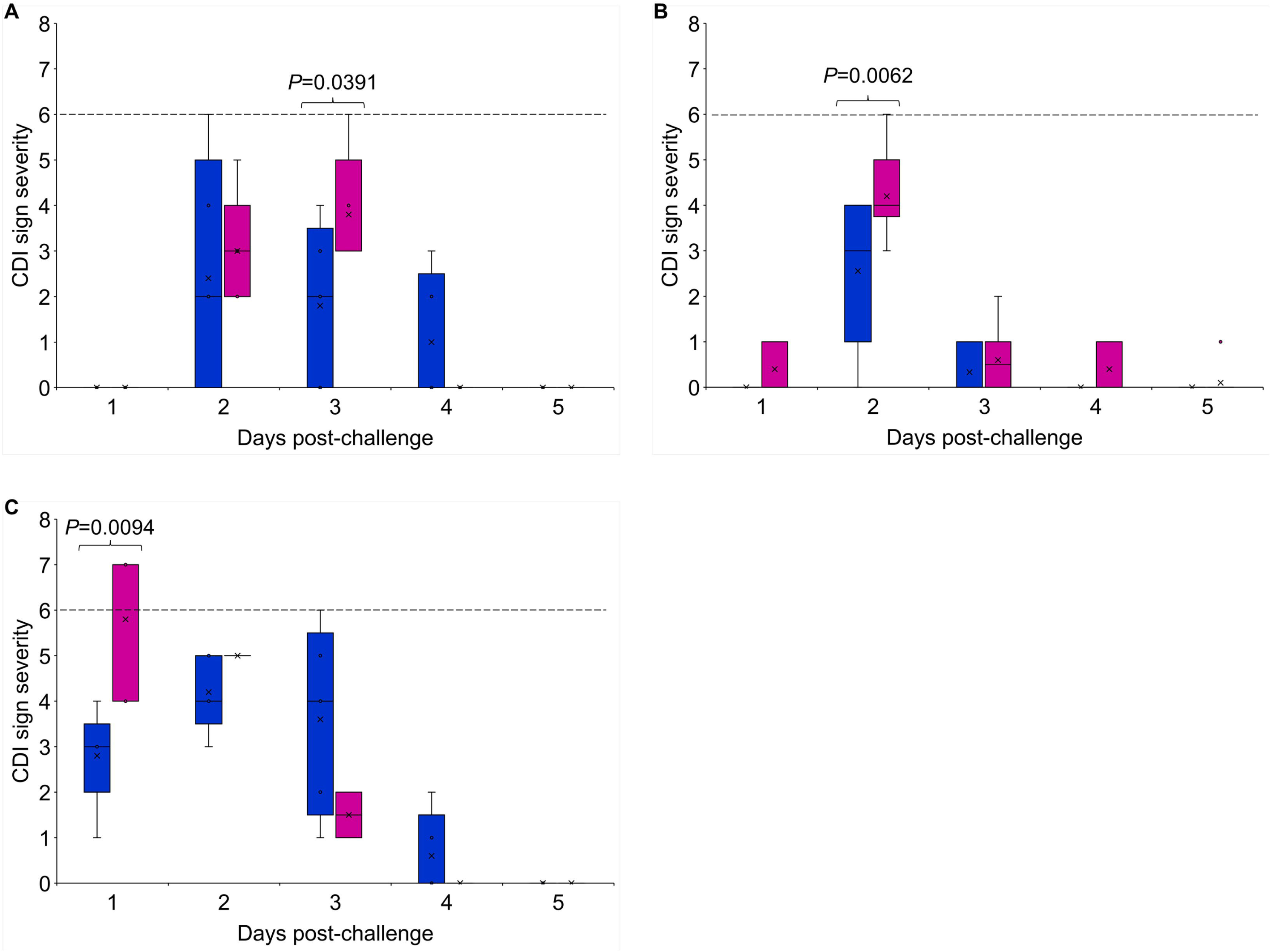
Effect of sex on CDI severity. Box-and-whisker plots of CDI sign severity in female mice (pink bars) and male mice (blue bars) challenged with *C. difficile* spores from (A) strain 9001966, (B) strain 630, and (C) strain R20291. Signs of severity were determined daily based on the CDI scoring rubric reported previously [36]. Animals with a score of <3 were indistinguishable from noninfected controls and were considered non-diseased. Animals with a score of 3 to 4 were considered to have mild CDI. Animals with a score of 5 to 6 were considered to have moderate CDI. Animals with a score of >6 were considered to have severe CDI and were immediately culled. Dashed line represents the clinical end point. Horizontal bold lines represent median values, asterisks represent mean values, bars represent interquartile ranges, error bars (whiskers) represent maximum and minimum values, and dots represent inner point values. Daily statistical differences between males and females were determined by one-tailed *t*-test.

A second group of male and female mice was infected with spores from *C. difficile* strain 630 (ribotype 012). Male mice infected with *C. difficile* strain 630 spores showed less individual variability in CDI symptomatology and shorter infection progression compared to animals infected with *C. difficile* strain 9001966. Male mice infected with *C. difficile* strain 630 did not show any signs on day 1 after challenge but started to show CDI signs by day 2. Female mice started to develop mild CDI signs one day after spore challenge (**Fig. 1B**). By day 2, female mice showed a notable increase in CDI sign severity, followed by a return to mild CDI signs by day 3 and complete remission by day 5 post-challenge. Similar to female mice, CDI signs for male mice started to resolve by day 3 and disappeared by day 5 post-challenge. In both male and female mice infected with *C. difficile* strain 630, CDI sign severity peaked on day 2 post-challenge. Notably, male mice showed mostly mild-to-moderate sign severity, while female mice showed statistically more severe symptomatology (**Fig. 1B**).

A third group of male and female mice was infected with spores from hypervirulent *C. difficile* strain R20291 (ribotype 027). Male mice challenged with *C. difficile* strain R20291 started to develop moderate CDI symptoms by day 1 post-spore challenge (**Fig. 1C**). Male mice remained moderately symptomatic until day 4, when CDI signs started to resolve. Similar to animals infected with *C. difficile* strain 9001966 and *C. difficile* strain 630, all male mice infected with *C. difficile* strain R20291 recovered and showed no CDI signs by day 4.

In contrast, female mice challenged with *C. difficile* strain R20291 spores developed statistically more severe CDI signs than male mice the day after spore challenge (**Fig. 1C**). Indeed, while all males infected with *C. difficile* strain R20291 recovered from CDI, more than half of the female cohort succumbed to the infection (**Fig. 2B**, p=0.049). Surviving female mice started to slowly recover from CDI and showed no symptoms by day 4. Because animals challenged with *C. difficile* strain R20291 developed the most severe CDI signs, we used this hypervirulent strain in subsequent experiments.

**Figure 2.**
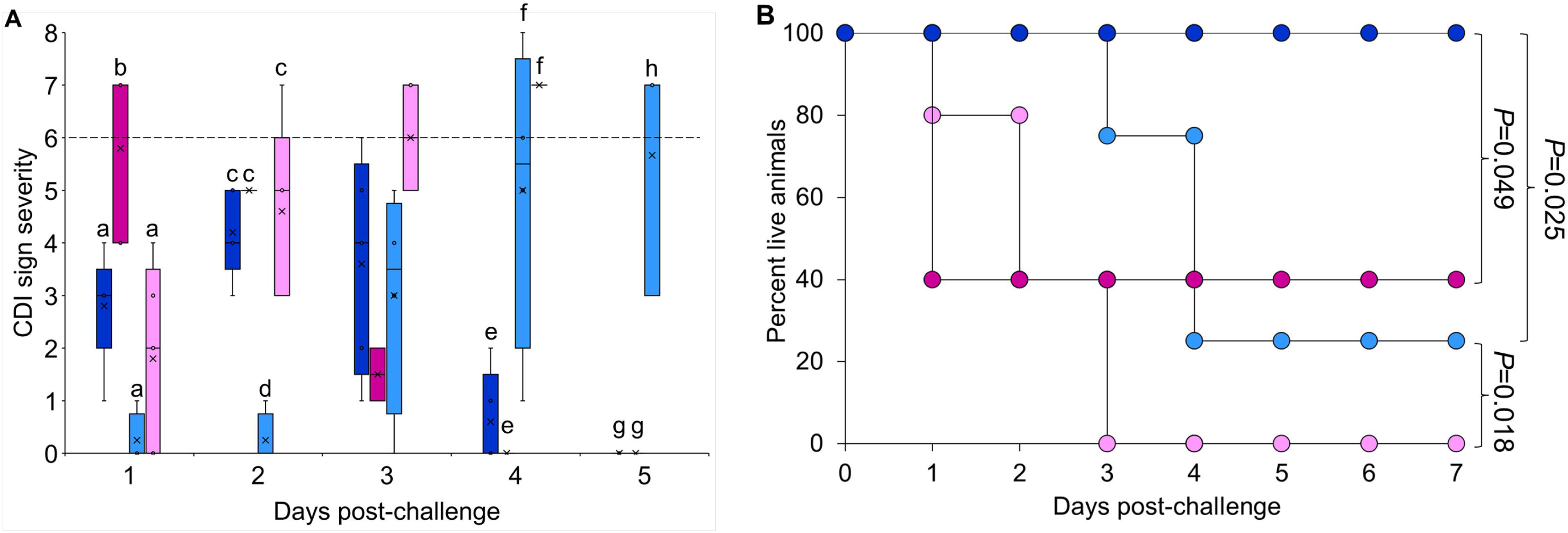
Effect of Atkins-style diet on mice CDI survival. (A) Box-and-whisker plots of CDI sign severity and (B) Kaplan-Meier survival plots for male mice fed a standard diet (solid blue), female mice fed a standard diet (solid pink), male mice fed an Atkins-type diet (blue crosshatch), and female mice fed an Atkins-type diet (pink crosshatch) challenged with *C. difficile* spores from strain R20291 [36]. For panel A, horizontal bold lines represent median values, asterisks represent mean values, bars represent interquartile ranges, error bars (whiskers) represent maximum and minimum values, and dots represent inner point values. Single-factor ANOVA was performed at every time point to assess differences among the sign severity means of the different groups. ANOVA results with *P* values of <0.05 were analyzed *post hoc* using the Scheffe’s test for pairwise comparison between daily groups. Columns that are labeled with different letters are statistically different. For panel B, statistical survival comparisons of different groups were performed via log rank test.

### A high-protein/high fat (Atkins-type) diet aggravates CDI in mice of both sexes

We recently showed that an Atkins-type diet can exacerbate CDI severity in mice [36]. To determine if sex affects CDI exacerbation by diet, male and female mice challenged with *C. difficile* strain R20291 while on this high protein/high fat (Atkins-type) diet showed delayed onset of CDI signs compared to animals fed a standard diet (**Fig. 2A**). Although female mice fed an Atkins-type diet did not show statistically more severe CDI signs (**Fig. 2A**) or survival (**Fig. 2B**) than female mice fed a standard diet, by day 4 post-challenge all female mice had developed severe CDI and had to be culled.

In contrast, once the infection started, male mice fed an Atkins-type diet developed statistically more severe CDI signs compared to challenged mice fed a standard diet (**Fig. 2A**). On a standard diet, 100% of males survived compared to 40% of females (p=0.049). Indeed, while all male mice on a standard diet survived CDI, 75% of male mice fed an Atkins-type diet developed signs severe enough to require euthanasia (**Fig. 2B**). In fact, survival for male mice on an Atkins-type diet was statistically worse than male mice fed a standard diet (p=0.025) comparable to female mice fed a standard diet, but still better than females on the Atkins-type diet (p=0.018).

### Estrous stages have a delayed effect on CDI severity prior to infection

Using vaginal smear cytology (**Figure S1**), we were able to monitor the estrous cycle of female mice. Animals were binned (**Fig. 3**) into estrus stage (blue columns), metestrus stage (orange columns), diestrus stage (green columns), and proestrus stage (magenta columns). Some animals could not be unambiguously assigned to a specific estrous stage. These animals showed borderline cytology readings and were thus binned into two new categories: late diestrus/early proestrus (pink columns) and late proestrus/early estrus (yellow columns).

**Figure 3.**
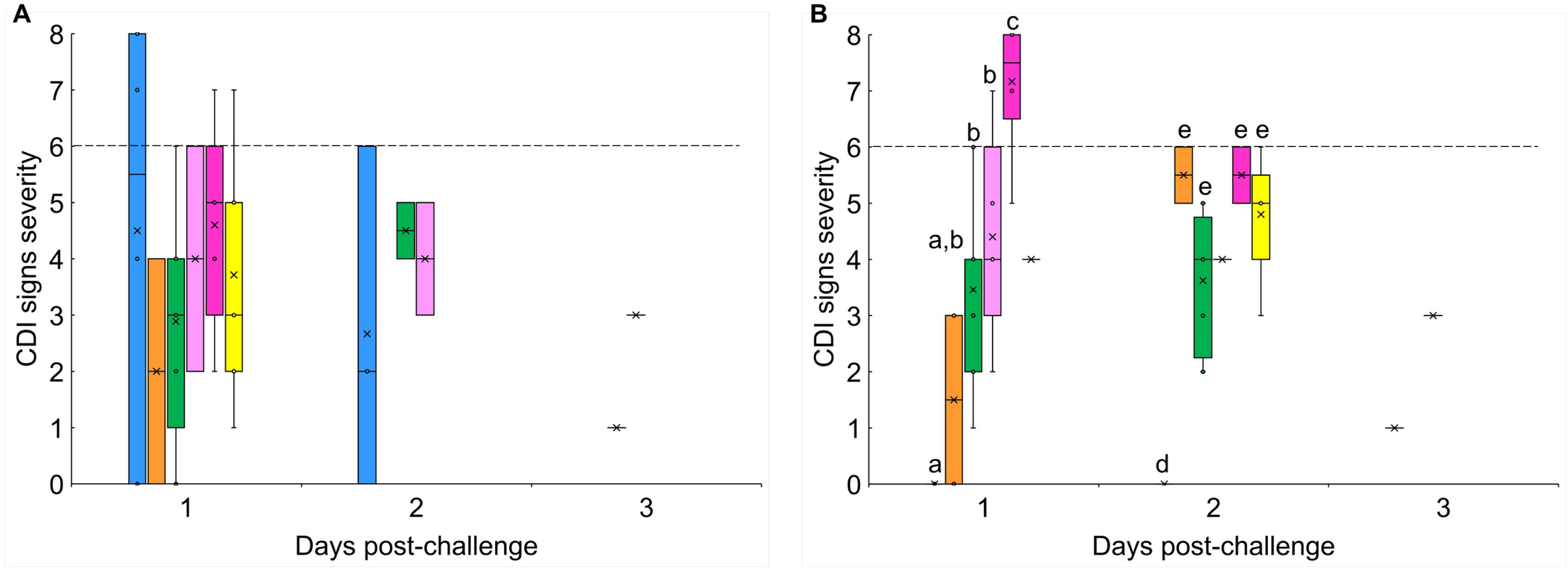
Time-dependent effect of estrous cycle on daily CDI severity. Box-and-whisker plots of daily CDI sign severity plotted against (A) their concurrent estrous stages and (B) their prior-day estrous stages. Animals in estrus (blue bars), metestrus (orange bars), diestrus (green bars), late diestrus/early proestrus (pink bars), proestrus (magenta bars), and late proestrus/early estrus (yellow bars) were challenged with *C. difficile* spores from strain R20291 on day 0. CDI sign severity and estrous cycle stages were determined daily. Signs of severity were determined based on the CDI scoring rubric reported previously [36]. Horizontal bold lines represent median values, asterisks represent mean values, bars represent interquartile ranges, error bars (whiskers) represent maximum and minimum values, and dots represent inner point values. Single-factor ANOVA was performed at every time point to assess differences among the sign severity means of the different estrous stages. ANOVA results with *P* values of <0.05 were analyzed *post hoc* using the Scheffe’s test for pairwise comparison between daily groups. Columns that are labeled with different letters are statistically different.

To determine the effect of the estrous cycle on murine CDI severity, a first set of female mice in distinct stages of the estrous cycle were challenged with *C. difficile* strain R20291 spores. We continued to simultaneously monitor the estrous cycle and CDI sign progressions. As the estrous cycle does not stop during the course of infection, animals could cycle across different estrous stages while CDI signs are being monitored. As a first approach, we tried to correlate daily CDI signs for each animal with their concurrent estrous stage, but we found no statistical significance between groups (**Fig. 3A**).

Intriguingly, when daily CDI severity was plotted against the estrous stages of the prior day, we found distinct daily patterns (**Fig. 3B**). The strongest statistical correlation between CDI signs and prior day estrous stages was found on day 1 post-challenge. Animals that were in estrus when challenged with *C. difficile* spores (day 0), showed no CDI signs the following day (day 1). Compared to animals that were in estrus on day 0, animals that were in metestrus, diestrus, and late diestrus/early proestrus during spore challenge developed statistically higher CDI severity the following day. Finally, animals that were at proestrus at day 0 developed the most severe CDI symptoms the day after challenge with more than half reaching the clinical endpoint.

The effect of prior-day estrous stages on CDI sign severity started to dissipate by day 2 post-challenge. On day 2 post-challenge, animals that were in estrus on day 1 showed no CDI symptoms. In contrast, animals that were in metestrus, diestrus, or late proestrus/early estrus on day 1 post-challenge showed statistically higher CDI sign severity. Unfortunately, there was an insufficient number of animals in the proestrus and the late diestrus/early proestrus groups on day 1 for a complete statistical analysis of CD severity on day 2 post-challenge. Furthermore, because of the high CDI-mediated mortality on days 1 and 2, there were insufficient numbers of animals left for statistical analysis on CDI score on day 3 post-challenge vs estrous stages.

To determine if the prior-day estrous effect is independent of the infection time course, we combined the data for each estrous stage regardless of the day they were determined. This allowed us to increase the number of animals in each estrous stage for statistical analysis. As expected, plotting CDI severity against their concurrent estrous stages showed no statistically significant differences (**Fig. 4A**).

**Figure 4.**
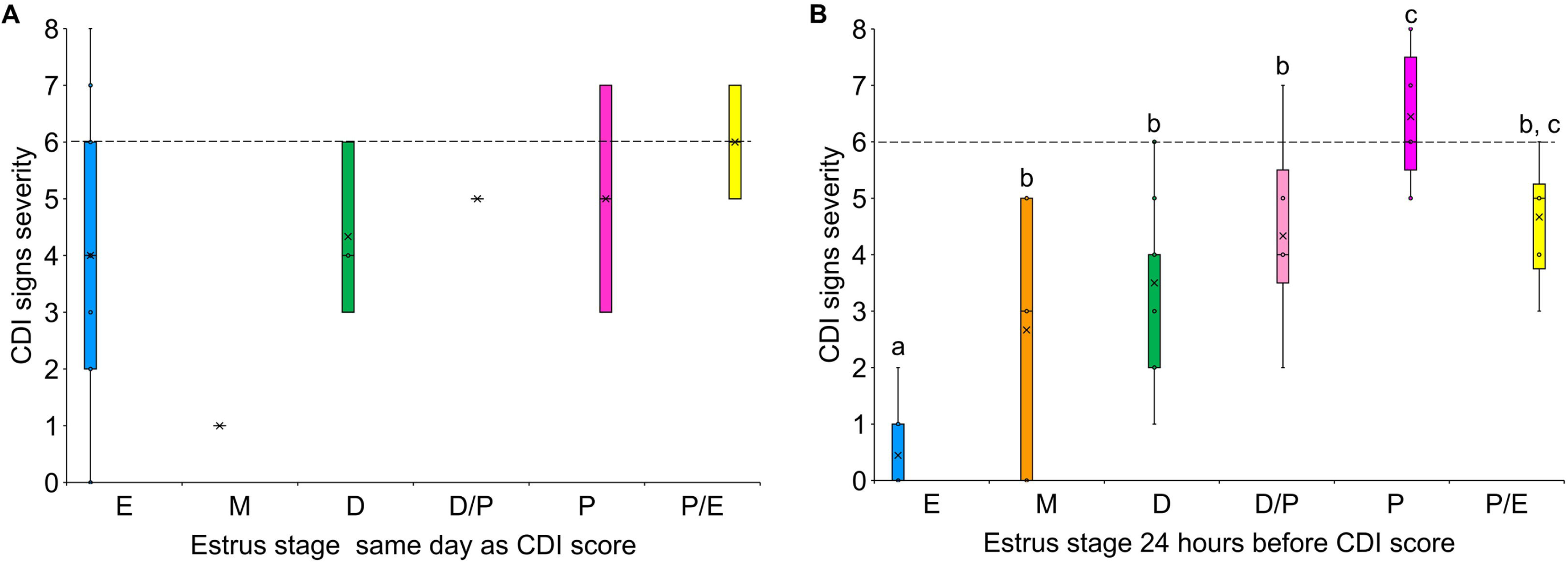
Time-independent effect of estrous cycle on CDI severity. Cumulative box-and-whisker plots of CDI sign severity caused by *C. difficile* strain R20291 plotted against (A) their concurrent estrous stages, and (B) their prior-day estrous stages. Animals that were at estrus (blue bars), metestrus (orange bars), diestrus (green bars), late diestrus/early proestrus (pink bars), proestrus (magenta bars), and late proestrus/early estrus (yellow bars) at any day were pooled together. CDI sign severity and estrous cycle stages were determined daily. Signs of severity were determined based on the CDI scoring rubric reported previously [36]. Horizontal bold lines represent median values, asterisks represent mean values, bars represent interquartile ranges, error bars (whiskers) represent maximum and minimum values, and dots represent inner point values. Single-factor ANOVA was performed to assess differences among the sign severity means of the different estrous stages. ANOVA results with *P* values of <0.05 were analyzed *post hoc* using the Schiffer test for pairwise comparison between all groups. Columns that are labeled with different letters are statistically different.

However, when CDI severity for each animal was plotted against their prior-day estrous stage, a similar delayed pattern was observed as in **Fig. 3B**. Indeed, when animals were in estrus at any day during infection, they showed few CDI signs the next day (**Fig. 4B**). Animals that were in metestrus, diestrus, and diestrus/proestrus at any time showed more varied but similar intermediate symptomatology one day later. In contrast, animals in proestrus at any time during infection became extremely sick the next day and had to be culled. Any animals in proestrus/estrus showed next-day CDI severity intermediate between proestrus and estrus.

### Estrous stages are correlated with day of maximal CDI sign severity

During the previous part of the study, only vaginal lavage samples were collected and examined to determine the estrous stage of animals. To further investigate the delayed effect of the estrous stages on CDI severity, a second cohort of mice were tracked and timed for blood collection and sacrificed based on their predicted estrous stage on the day of spore challenge with hypervirulent strain R20291. Instead of collecting terminal blood samples, facial vein blood samples were taken 1-2 days prior to sacrifice. Contrary to expectations, mice did not develop maximal CDI signs 1-day post-challenge as the previous cohort of mice (**Fig. 1A**). Instead, all mice developed the most severe CDI signs 2 days after infection. Consequently, no correlation was observed between estrous stage (via vaginal lavage and cytology) and day 1 post-challenge.

Since the maximum CDI signs for the second cohort did not develop until 2-days post-challenge, the temporal effect of the estrous stage was also delayed (**Fig. 5**) compared to animals in the first cohort (**Fig. 4B**). Indeed, second cohort animals that were in the estrus stage showed mild CDI signs two days later. In contrast, second cohort animals in the proestrus stage developed more severe CDI signs two days later.

**Figure 5.**
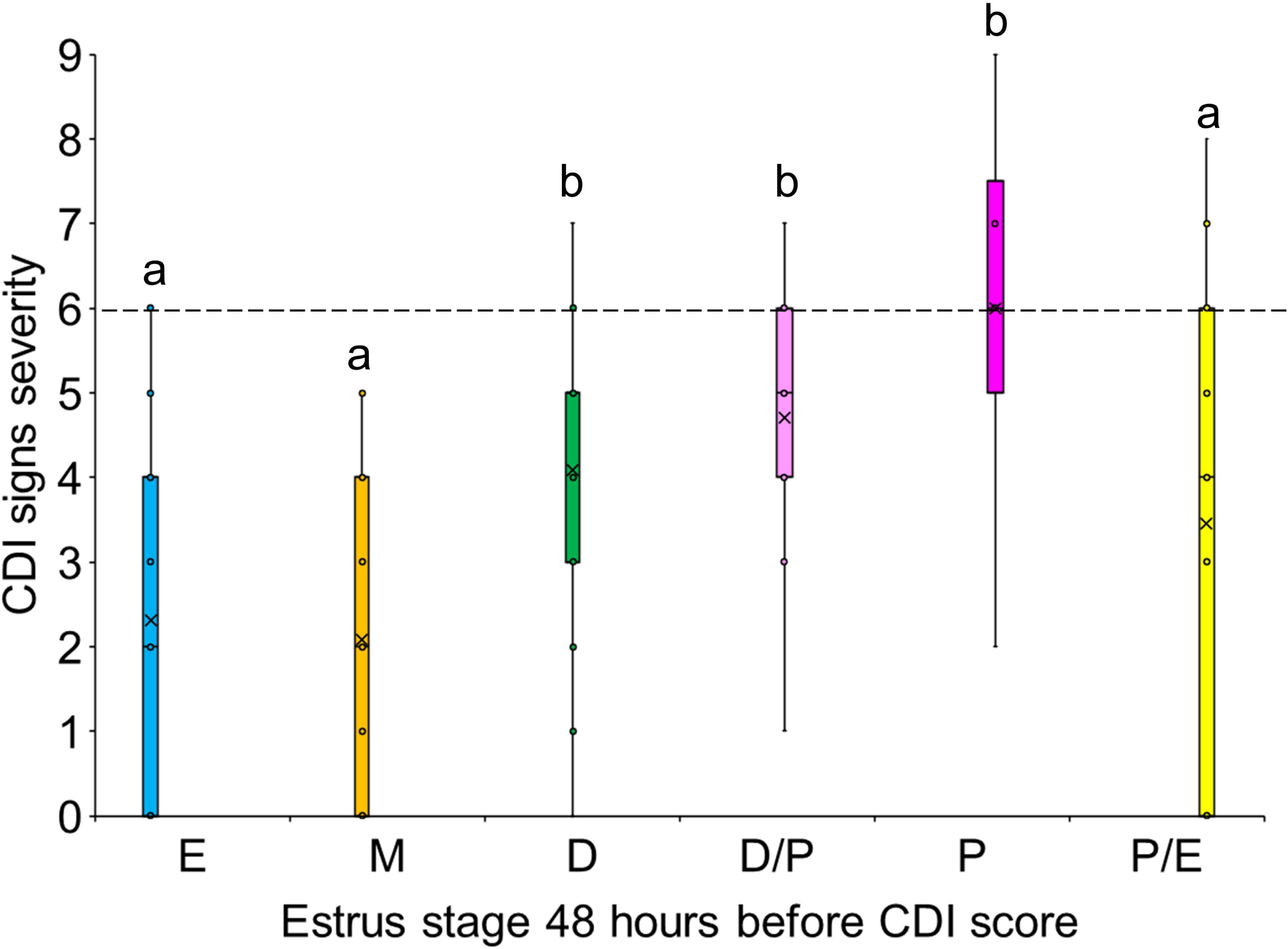
CDI severity score and estrous stage 48 prior. were scored daily for 5-7 days following infection, and estrous stages were simultaneously tracked daily by microscopic examination of cells collected by vaginal lavage. The maximum CDI score for each mouse was determined and compared to its estrous stage 48 hours prior to the maximum score. Single-factor ANOVA was performed to assess differences among the sign severity means of the different estrous stages. ANOVA results with *P* values of <0.05 were analyzed *post hoc* using the Schiffer test for pairwise comparison between all groups. Columns that are labeled with different letters are statistically different.

### Biomarkers obtained before infection (D0) and during early infection (D1)

Pre-infection blood samples (D0, obtained hours before spore challenge) and early infection blood samples (D1, one day after spore challenge) were obtained from female mice to test for the effect of hormones, cytokines, and immunoglobulins on CDI progression. As expected, pre-infection (D0) hormone levels varied between individuals since animals were at different estrous stages when they were challenged with *C. difficile* spores on day 0. Interestingly, we also found inter-individual variabilities for pre-infection levels of cytokines and immunoglobulins, even though blood was collected before spore challenge. In contrast, early infection (D1) cytokine and immunoglobulin levels were more varied, as expected for animals progressing through active infection and showing different sign severities.

## Discussion

Few attempts have been made to understand how sex affects CDI onset and severity. To assess the clinical importance of sex on human CDI, we did an extensive PUBMED search for primary articles relating to CDI risks. We found that most studies (n=87) did not stratify for sex. Of the studies that assessed sex as a variable (n=24), most linked CDI to other comorbidities.

These studies tended to have small sample sizes, making it difficult to determine if sex was associated with CDI or rather to the underlying comorbidity. However, when CDI was the primary diagnosis, multiple large cohort studies showed that women are at higher risk than men of developing symptoms [7, 8, 10, 11, 47]. Indeed, a multicenter retrospective cohort study of over 1 million patients at 150 US hospitals showed that the odds of CDI in women was 1.2 times as likely as in men [47]. Although the correlation between sex and human CDI is clear, testing for susceptibility in women is cost prohibitive and impractical. Yet patient outcomes could be dramatically improved if clinicians could adapt antibiotic treatments to match specific hormone levels.

In this study, the murine model recapitulates many of the differences in CDI outcome observed between men and women [7, 8, 10, 11, 47]. For instances, during our investigations into the factors affecting CDI progression and prophylaxis [33, 40, 41], we serendipitously found that female mice developed more severe CDI signs than males.

To further test for sexual dimorphism in the murine model of CDI, we infected mice with three different *C. difficile* strains. These strains represent different ribotypes and were selected because they show distinct timelines for CDI onset. *C. difficile* strain 9001966 (ribotype 106) is a clinical strain from the United Kingdom and shows delayed onset of murine CDI. *C. difficile* strain 630 (ribotype 012) is a type strain [48, 49] that we have extensively tested for its capacity to induce CDI symptoms in mice [40, 41]. Finally, *C. difficile* strain R20291 (ribotype 027) is a hypervirulent strain that causes earlier infection onset and more severe murine CDI signs than other strains [33].

When animals were infected with any of these strains, female mice developed statistically more severe CDI than their male counterpart. The timing of these differences varied, but in every case, the most significant differences in CDI severity coincided with the peak of disease severity in their respective infection timeline. Because animals challenged with hypervirulent *C. difficile* strain R20291 showed the strongest CDI signs, we continued to use this strain for all subsequent experiments.

In contrast to hamsters [38, 50], murine CDI seldom tends to be lethal [33, 40, 41]. However, we recently showed that an Atkins-type (high-protein/high-fat) diet can greatly increase murine CDI sign severity duration and mortality [36]. Here, we determined if the Atkins-type diet affected CDI severity between male and female mice. When animals were fed a diet known to exacerbate CDI signs, mortality of male mice increased significantly. Even then, male mice showed less severe signs than females fed the same diet. This shows that the sexual dimorphism of murine CDI is still present under conditions where CDI is severe.

The most obvious difference between male and female mice is the female estrous cycle. However, intraday correlation between estrous cycle and CDI severity showed no statistical differences at any time during the infection. In contrast, estrous stages seem to have a 1 day to 2 days delayed effect on CDI severity.

The interplay between estrous stage cycling and CDI progression is quite complex. For CDI caused by *C. difficile* strain R20291, infection starts and reaches a maximum of severity on days 1 or 2 following spore challenge. Surviving animals completely recover by days 4-5 post-challenge. In contrast, the estrous cycle can take between 3-6 days to be completed. Furthermore, the duration of each estrous stage is variable, with proestrus and metestrus lasting as little as 8 hours and diestrus lasting as long as 72 hours [51]. This was further complicated by the observation that estrous stages showed a 1–2-day delayed effect on CDI severity. Thus, binning animals into different estrous stages for each day reduced our statistical power for some of the estrous stages. Nevertheless, our data clearly shows that animals challenged while in the estrus stage were largely protected from CDI. In contrast, animals challenged while in the proestrus stage developed deadly CDI.

The contrast between proestrus-mediated and estrus-mediated CDI susceptibility could not be starker. While CDI is most deadly following proestrus, estrus offers strong CDI protection to animals on the typical day of maximal disease severity whether1 day or 2 days after estrous staging. Furthermore, metestrus and diestrus correlate with more heterogeneous CDI symptomatology on days of maximal onset.

An obvious factor underlying CDI sexual dimorphism is the fluctuating hormone levels during the female ovulation cycle [19]. In general, CDI-exacerbating proestrus is characterized by large increases in estradiol, progesterone, LH, and FSH levels and low concentrations of prolactin [29]. These spikes in hormone levels signal entry into the CDI-protective estrus. At this point, the high concentrations of circulating estradiol, progesterone, LH, and FSH characteristic of proestrus are severely diminished followed by an increase of serum PRL. The metestrus and diestrus stages tend to have hormonal levels that fall in between those of proestrus and estrus [29].

Correlation analysis of pre-infection (D0) biomarkers form two, almost separate cluster. In fact, cytokine KC is the only connection between the two clusters. Cluster 1 (**Fig. 6**, top) is centered around pituitary hormones FSH, PRL, and LH. Cluster 2 (**Fig. 6**, bottom), on the other hand, is centered around steroidal hormone progesterone.

**Figure 6.**
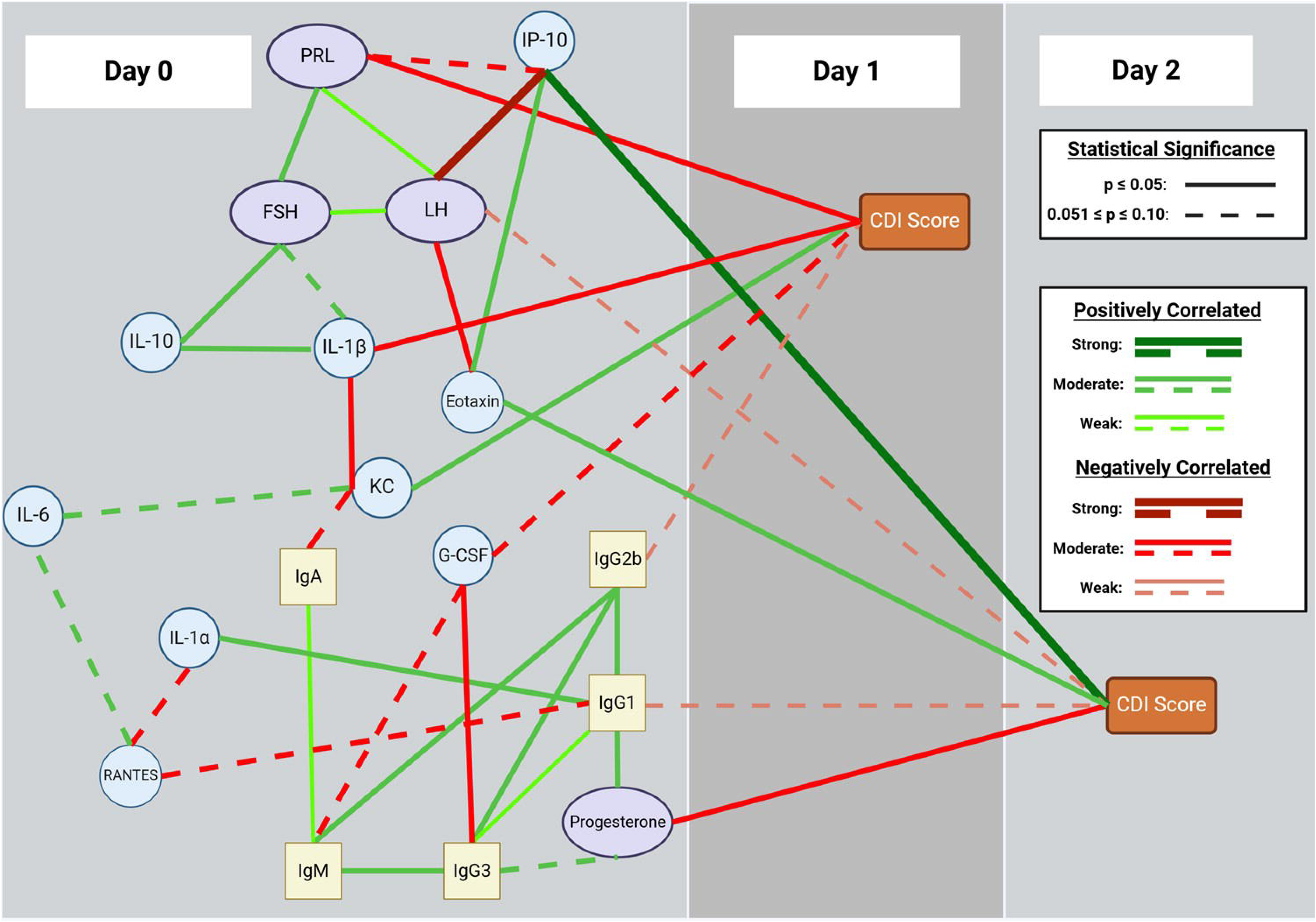
Biomarker networks before infection. Serum was obtained from female mice before challenge (D0) with *C. difficile* spores and individually analyzed for concentrations of sex hormones, cytokines, and immunoglobulins. Animals were scored for CDI sign severity on days 1-5 post-challenge. Days with correlations between biomarkers and CDI severity are highlighted (days 1 and 2 post-challenge). Biomarker levels and CDI scores were correlated using Kendall-tau. Positive correlations are represented with green lines. Negative correlations are represented with red lines. Line thickness and hues represent the strengths of the correlations. Solid lines represent statistically significant correlations with p ≤ 0.05. Dashed lines represent statistically significant correlations with 0.051 ≤ p ≤ 0.10.

From cluster 1, pre-infection (D0) levels of hormone LH, hormone FSH, and cytokine IL-10 form a positively correlated network that negatively influences CDI severity during the following day (day 1 after spore challenge) through PRL and cytokine IL-1β. Only cytokine KC positively correlate with CDI sign on day 1 post-challenge while simultaneously negatively correlating with IL-1β and IgA levels and positively correlating with IL-6.

From cluster 2, pre-infection (D0) levels of hormone progesterone, cytokine IL-1α, immunoglobulin IgG1, immunoglobulin IgG3, immunoglobulin IgM, and immunoglobulin IgG2b form a complex positively correlated group that negatively affect CDI severity on the following day through immunoglobulin IgG2b. From the same group, IgM and IgG3 negatively correlate with cytokine G-CSF. In turn, G-CSF negatively influences CDI score on day 1.

Mirroring the temporal delay between estrous stages and CDI severity, pre infection (D0) levels of PRL, FSH, LH, cytokine IP-10, and cytokine eotaxin form a group that positively correlate with CDI severity two days post-challenge. The same network negatively affect CDI severity two days post-challenge trough LH The two-day delayed effect is also observed in cluster 2. Pre infection (D0) levels of the progesterone/immunoglobulin group negatively influences CDI severity two days post-challenge through IgG1 and progesterone. As expected after spore challenge, early infection (D1) immunoglobulin, cytokine, and chemokine levels create a single complex network that correlate CDI severity on the same day (**Fig. 7**). Early infection levels of LH and FSH positively influence same day CDI sign severity through IFNγ and KC, respectively.

**Figure 7.**
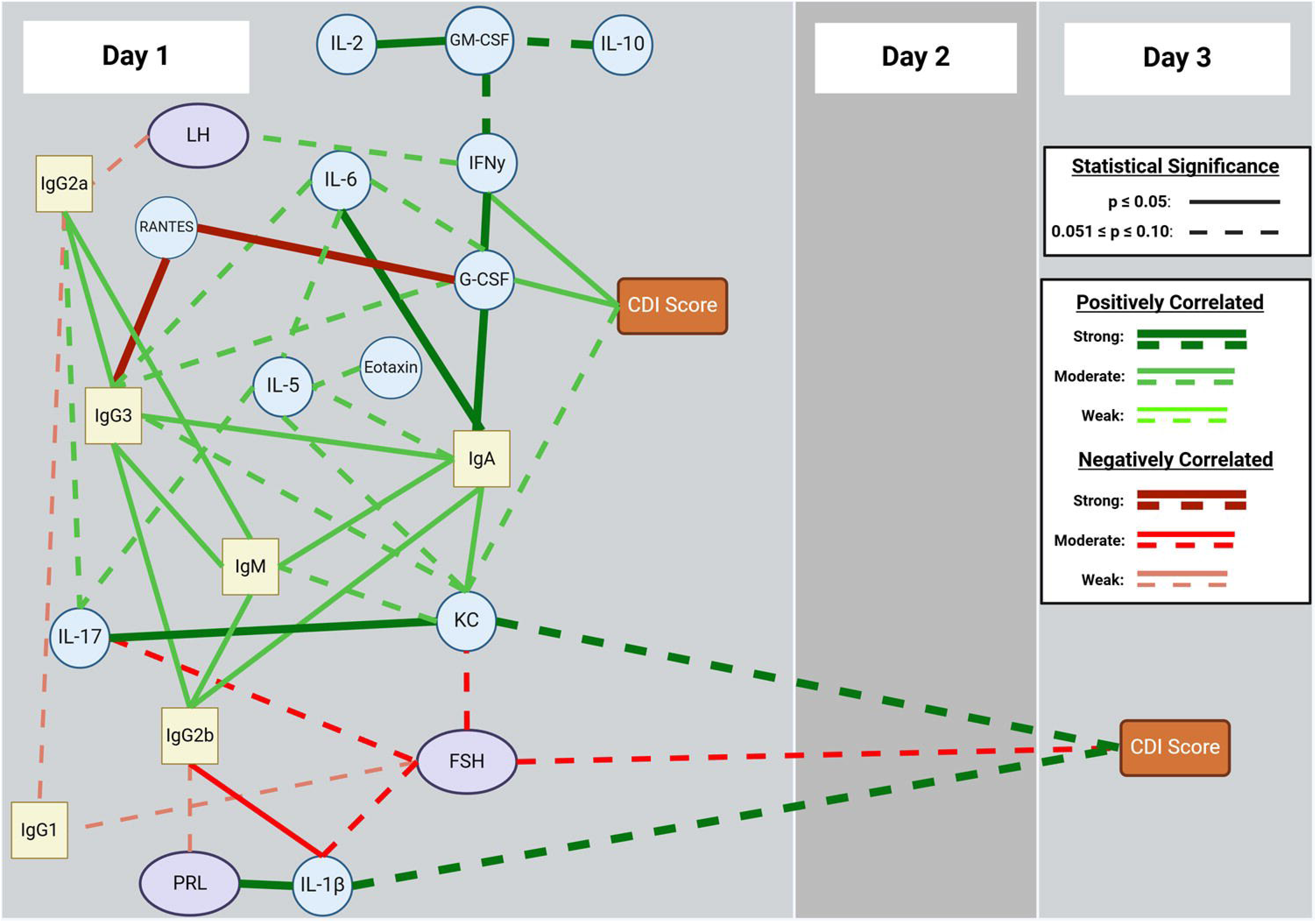
Biomarker networks during early infection. Serum was obtained from female mice the day after challenge (D1) with *C. difficile* spores and individually analyzed for concentrations of sex hormones, cytokines, and immunoglobulins. Animals were scored for CDI sign severity on days 1-5 post-challenge. Days with correlations between biomarkers and CDI severity are highlighted (days 1 and 3 post-challenge). Biomarker levels and CDI scores were correlated using Kendall-tau approaches. Positive correlations are represented with green lines. Negative correlations are represented with red lines. Line thickness and hues represent the strengths of the correlations. Solid lines represent statistically significant correlations with p ≤ 0.05. Dashed lines represent statistically significant correlations with 0.051 ≤ p ≤ 0.10.

We also observed a temporal effect between early infection biomarkers and CDI recovery, two days later. Indeed, early infection (D1) levels of KC and IL-1β show strong positive correlation to CDI severity on day 3 post-challenge. Central to these effects, early infection (D1) FSH levels shows strong negative correlation with early infection (D1) levels of KC and IL-1β. FSH also shows negative correlation with CDI severity on day 3 post-challenge.

We have previously shown that *C. difficile* spores germinate between 6- and 9-hours post-challenge in the intestines of mice. Potentially, the strong association between day 0 estrous stages, pre-infection biomarker levels, and CDI severity on days 1-2 could be due to estrous cycle and hormone fluctuations influencing the initial stages of CDI onset. On the other hand, delayed effects of the estrous cycle and sex hormones seem to continue throughout CDI progression into infection recovery. This suggests that estrous stages can increase or decrease female mice susceptibility to CDI both at the early and late stages of the infection course.

The negative correlation of pre-infection PRL (D0) with early CDI signs mirrors the protective effect of the estrus cycle. However, the fact that pre-infection (D0) levels of progesterone and LH show a time-delayed, negative correlation to CDI severity is more difficult to explain. It should be also noted that the boundaries between estrous stages are arbitrary and dependent on cellular response to hormonal changes. In contrast, changes in sex hormone levels follow a time continuum throughout the estrous cycle. For example, even though progesterone is expected to fall during estrus, entry into the stage happens when progesterone levels are at its maximum. Hence, small variations in sample collection timing, could result in uncoupling between expected sex hormone levels and identified estrous stages. Indeed, murine sex hormone fluctuations have been shown to follow strain-specific patterns (REF) and even show significant variabilities between individuals on the same assigned estrous stages. These temporal uncertainties could partially explain the paradoxical nature of sex hormone effects on CDI severity.

### Conclusions

Human CDI is not diagnosed unless symptoms are severe enough to warrant clinical intervention. In that sense, CDI diagnosis serves as a surrogate for symptoms severity in humans. Similar to other infection models, murine CDI provides an idealized and controlled (albeit imperfect) model to study disease onset, severity, and progression. The murine CDI model presents symptoms progression similar to human CDI and animals respond to the same treatments [52, 53]. We have found that murine CDI follows a well-ordered progression of disease signs. Based on these observations, we have implemented a rubric for CDI severity development [33, 41].

Little is known regarding sex hormones, their role in intestinal infection, and the adaptation of the immune response in the GI tract. The delayed effect of the estrous stage and sex hormones on CDI severity might be due to the time needed for these hormones to affect host physiological (e.g., immune system, intestinal microbiota, steroidal hormone accumulation in the gut) responses during infection.

During pregnancy in humans and mice, progesterone stimulates the expression of IL-4 and IL-10 and favors the activation of Th2 cytokines while suppressing the major Th1 cytokines, IL-12p70, IL-2, IFN-y, IL-18 [54–56]. However, little is known about the role of progesterone in intestinal infections. A recent study showed that exposure to progesterone resulted in decreased *Listeria monocytogenes* burden in cultured intestinal cells [57].

PRL is both a hormone and a cytokine with roles in reproduction, lactation, and stress regulation [58]. PRL has been shown to have immune modulating effects. In this study, pre-infection PRL levels affect early CDI progression. This is consistent with typical chronological presentation of hormones in the murine estrous cycle with PRL levels rising approximately one day (estrus stage) prior to elevation of progesterone levels (metestrus-diestrus stage). However, pre-infection PRL does not seem to correlate with any cytokine, chemokine, or immunoglobulin that that affect early CDI progression. Thus, the mechanism by which PRL affect CDI onset is not clear.

Of all the immunoregulatory compounds tested, pre-infection PRL only correlated with IP-10 levels. Similar to what we observed in this murine model, a strong direct correlation between severe CDI and upregulation of IP-10 has been observed in humans [59]. Hence, pre-infection PRL spike during murine protective-estrus stage, may cause a decrease in pre-infection IP-10 production resulting in less severe disease two days later. In our study progesterone did not show any significant direct relationships with any chemokines and cytokines. Nevertheless, animals with higher levels of pre-infection progesterone were protected from CDI onset two days later. In contrast to the lack of effect on cytokines, pre-infection progesterone forms a positive correlated group with almost all pre-infection immunoglobulins tested. Although progesterone’s effect on immunoglobulins is not well understood, progesterone’s immunomodulatory capabilities suggest that it can potentially influence immunoglobulin productions. Indeed, murine, human, and nonhuman primate, cervicovaginal IgG subtypes are highest during times of increased progesterone [60].

We observed a similar positive correlation between pre-infection progesterone levels and pre-infection IgG subtypes. In turn, both IgG1 and IgG3 showed correlations with cytokines and chemokines. IgG1 positively correlated with IL-1a, a dual role cytokine involved in homeostasis and the immune response [61] and negatively correlated with proinflammatory cytokine RANTES. In turn, IgG3 negatively correlated with proinflammatory cytokines G-CSF and RANTES.

Stimulated macrophages and other immune cells produce G-CSF and RANTES in response to infection. Both are important as chemoattractants and aid in maturation and activation of immune cells [62, 63] In fact, *C. difficile* has been shown to initiate a strong G-CSF and RANTES expression in the mice intestinal tract which may be used as markers for severity of disease [64, 65]. The direct and indirect effects of progesterone on immunoglobulins and cytokines could account for protection from CDI in mice.

Previous work in the Abel-Santos laboratory has shown that progesterone can inhibit *C. difficile* spore germination *in vitro*. Interestingly, sex steroidal hormones are also present at low concentrations in the intestinal tract and show a 1-day accumulation delay compared to serum [32], a timeline reminiscent to the delayed effect of progesterone on murine CDI. Due to the heterogenous environment of the intestinal lumen, there could be localized areas of higher hormone concentrations where they could interact with *C. difficile* spores. Whether progesterone affects CDI indirectly by modulating the immune system or directly by inhabiting spore germination, needs to be elucidated.

The gonadotropin LH is typically regarded as a reproductive hormone but studies have shown a correlation to intestinal cells where LH binds to receptors expressed in the intestine and slows gastric motility [66]. To our knowledge, the role of LH in intestinal infections has not been reported. Similar to progesterone, we found a negative correlation between pre-infection LH levels and CDI severity two days later. Pre-infection LH formed a positive correlated group with the other two pituitary hormones, PRL and FSH. Furthermore, like PRL, LH also negative correlated with pre-infection IP-10 while also positively correlating with eotaxin. Pre-infection IP-10 and eotaxin levels strongly positively correlate to CDI severity two days later. IP-10 has been reported as a potential biomarker for severity of disease including CDI [59, 67]. Eotaxin is a chemokine that recruits eosinophils to areas of inflammation and has been shown to play a role in the onset of the murine estrus cycle [68]. similar to IP-10, higher eotaxin levels have been associated with more severe CDI in humans [69]. Eotaxin eosinophil recruitment has also been shown to be necessary to prevent severe tissue damage in CDI [69].

After infection onset, the effect of sex hormones on disease progression seems to weaken. This might be due to the complex immune response that occurs after infection onset. At this point, the robust feedback look of the immune system would become more important than the more subtly effects of sex hormones.

Nevertheless, we found a potential PRL/FSH effect on CDI recovery. Early infection PRL levels positively correlated with IL-1β. Meanwhile, early infection FSH negatively correlated with both IL-1β and KC. In a mutual feedback loop, IL-1β can stimulate the secretion of PRL and PRL can increase the secretion of IL-1β through stimulated differentiation of T cells [58, 70]. Although IL-1β is typically considered a pro-inflammatory signal stimulated by infection, IL-1β has been shown to be complex and also play a role in commensal microbiota regulation and maintenance [71]. We are likely seeing this early infection loop playing a role in CDI recovery. Little is known regarding the effect of FSH on cytokine and chemokine production in the intestine of humans or mice. In fact, receptors for FSH are not highly expressed in the intestine [72]. In osteoclast differentiation FSH has also been shown to induce the expression of IL-1β and IL-6 from monocytes [73, 74]. Hence, In the murine intestine, both pre-infection and early infection FSH may have a role in the upregulation of IL-1β and IL-10 to aid in tissue maintenance and repair. These complex interactions between hormones, immunoglobulins and cytokines/chemokines warrants more study to better understand the onset and progression of CDI in mice and humans. Approaching *C. difficile* infection by studying the hypothalamic-pituitary-gonadal (HPG) axis, allows us to determine how sex hormones and the estrus cycle affect the severity and progression of CDI. Many cells of the immune system contain receptors for sex hormones which can activate cytokines, showing a connection between immune modulation and sex hormones. The full host immune response initiated by infection with *C. difficile* spores is yet to be fully elucidated. Many reports have identified cytokine and chemokine correlations between expression and severity of disease with similarities and differences in these reports suggesting dual roles in pathogenicity and protection. There is likely a very complex crosstalk between the systemic host immune response, the local immune cells and intestinal microbiota resulting in a fine line between infection clearance and tissue damage.

Together, these data suggests that sex hormones could affect CDI symptomatology directly by modulating spore germination in the intestine and/or indirectly by globally changing the host’s gut microbiome and/or immune status. The effect of individual hormones on CDI severity needs to be established but provides an open door into explorations on the relationship between sex hormones, the gut, and CDI.

## Acknowledgments

This work was supported by the National Institute of Allergy and Infectious Diseases at the National Institute of Health [grants numbers R15AI164346, R16AI175022] to EAS, and by the National Institute of General Medical Sciences at the National Institute of Health [grant number P20GM12132] to NIPM.

