## Supplementary material for "Effects of sexual dimorphism and estrous cycle on murine *Clostridioides difficile* infection": Figure S1

**Supplementary materials**

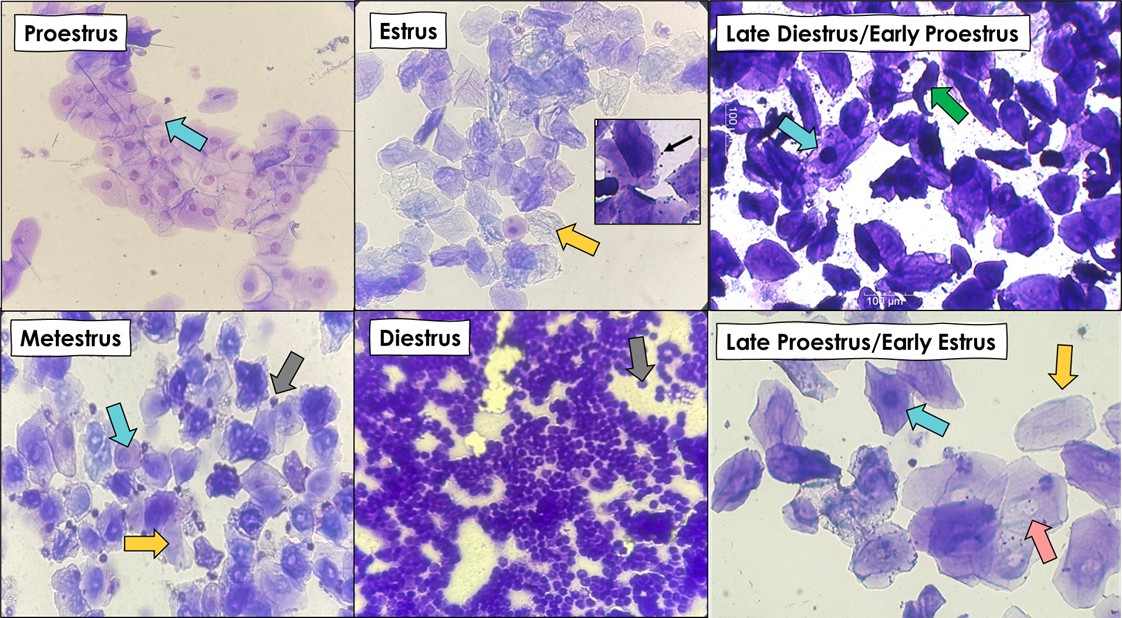

**Fig. S1.** **Estrous stages determination by vaginal cytology.** Vaginal swabs were taken and spread on microscopy glass slides. After air-drying, slides were stained with crystal violet. Estrous cycle stages can be identified based on the presence of either nucleated epithelial cells, cornified (keratinized) anucleated epithelial cells, leukocytes, or combinations of these cell types. The proestrus stage is dominated by the presence of nucleated epithelial cells. In contrast, when animals are in the estrus stage, cornified anucleated epithelial cells are in abundance. This stage can also show the presence of small specks of bacteria adhering to the epithelial cells. In the metestrus stage, all three major cell types are present. When animals transition to the diestrus stage, samples contain mostly leukocytes with scattered appearances of a few nucleated epithelial cells. Cytologic characteristics can also show intermediate stages of the estrous cycle. Small, irregular shaped cells can signify the formation of nucleated cells and are representative of late diestrus/early proestrus. Some lingering leukocytes may also be present at this stage. Late proestrus/early estrus is usually marked by a transition of nucleated cells to enucleation leaving a halo-like appearance in the middle of the cell.
